# FlopR: An open source software package for calibration and normalization of plate reader and flow cytometry data

**DOI:** 10.1101/2020.06.01.127084

**Authors:** Clare M. Robinson, Alex J. H. Fedorec, Ke Yan Wen, Chris P. Barnes

## Abstract

The measurement of gene expression using fluorescence markers has been a cornerstone of synthetic biology for the last two decades. However, the use of arbitrary units has limited the usefulness of this data for many quantitative purposes. Calibration of fluorescence measurements from flow cytometry and plate reader spectrophotometry has been implemented previously but the tools are disjointed. Here we pull together, and in some cases improve, extant methods into a single software tool, written as a package in the R statistical framework. The workflow is validated using *Escherichia coli* engineered to express GFP from a set of commonly used constitutive promoters. We then demonstrate its power by identifying the time evolution of distinct subpopulations of bacteria from bulk plate reader data, a task previously reliant on laborious flow cytometry experiments. Along with standardized parts and experimental methods, the development and dissemination of usable tools for quantitative measurement and data analysis will benefit the synthetic biology community by improving interoperability.

**Graphical Abstract:** 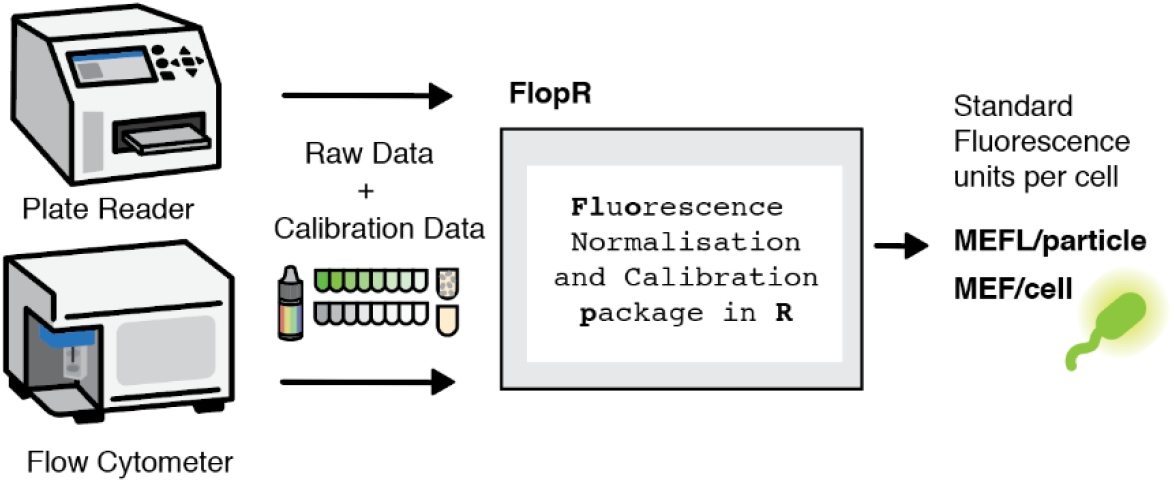

---

The construction of novel, reliable genetic circuits relies largely on the reproducible characterisation of standard genetic parts. Fluorescence is often used as a quantitative output for engineering genetic circuits and the characterisation of their individual parts and can be measured in microplate readers or flow cytometers. Plate readers are particularly suited for high throughput applications, such as understanding temporal dynamics of multiple different constructs and conditions simultaneously. However, the data are bulk measurements that obscure population heterogeneity. Flow cytometry reveals this information but has more limited throughput capabilities. Recently, an inexpensive and easily implementable protocol to convert green fluorescence measurements from both instruments into intercomparable standard units of molecules of equivalent fluorescein (MEFL) per particle was validated across hundreds of laboratories in the international Genetically Engineered Machine (iGEM) competition (1). Measurements of standard calibrants are used to create calibration curves to convert the arbitrary units of individual machines to standardized units, a technique which has previously been established (2–4). Currently, however, software tools for calibration and normalization exist across multiple platforms and different coding languages. Although this offers choice to end users, it makes high-throughput application difficult and time-consuming. In addition, autofluorescence normalization for plate reader measurements is often implemented inconsistently and in an ‘ad-hoc’ manner, even though it has been shown to be critical particularly when measuring fluorescent particles that share a similar emission wavelength with non-fluorescent cells (5). Consequently, standards for calibration and normalization have yet to be widely adopted and fluorescent data collected from microplate readers and flow cytometers is currently still often reported in arbitrary units.

Here we present FlopR, a full calibration and normalisation for flow cytometry and microplate reader data in R, the free and open source programming language. FlopR uses data from measurements of standard calibrants to generate and fit calibration curves and convert data recorded in arbitrary units to units of MEFL per particle. These calibrants are fluorescein and monodisperse silica microspheres for plate reader measurements and calibration particles for flow cytometry measurements. Fluorescein shares similar excitation and emission wavelengths to green fluorescent protein (GFP), and the microspheres used are of approximately the same size and diameter as *E. coli* (2). We demonstrate FlopR is able to accurately calibrate fluorescence data from *E. coli* that are constitutively expressing GFP consistently across multiple measurements on different days and in agreement with calibrated literature values. Finally, we show FlopR is able to reconstruct the growth of individual fluorescent subpopulations of cells from plate reader data of heterogeneous bulk cell cultures, demonstrating its potential for complex and high throughput applications such as studies of dynamics within mixed microbial consortia.

## Results

### Calibration Workflow

The full data processing workflow requires measurement of the fluorescent cells of interest, non-fluorescent negative control cells and calibrants specific to both machines, as shown in Figure 1. For the microplate reader calibration, conversion factors are generated from fluorescein and microsphere calibrants measured prior to the main experiment, and at least one well of blank media and non-fluorescent cells each measured during the experiment. For flow cytometry calibration, calibration beads and non-fluorescent cells are measured during the experiment with the same instrument settings. The wavelengths of the excitation laser and detection bandpass filter of both instruments must correspond for their final calibrated measurements to be comparable.

**Figure 1.**
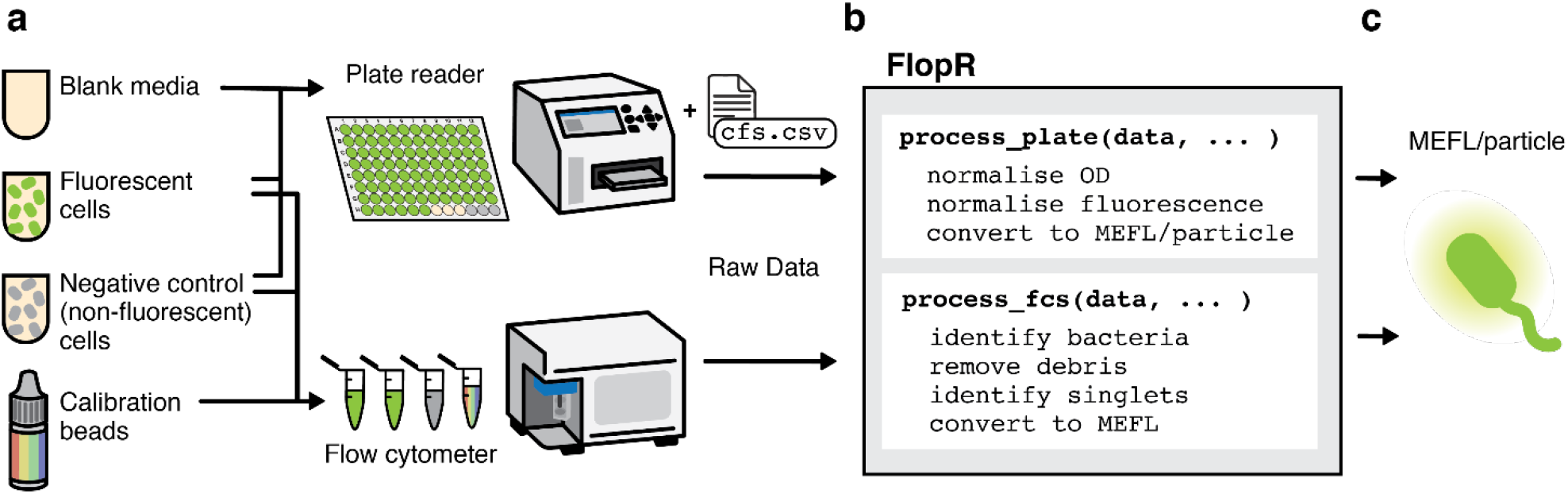
FLOPR work flow. (a) FlopR takes as input parsed, raw data from plate readers or flow cytometer, along with calibrants: non fluorescent cells and blank media (for plate reader) or calibration beads (for flow cytometry). (b) Data is processed by the FlopR package. (c) The output is processed data in comparable units of MEFL per particle independent of instrument.

To process plate reader data:

- absorbance of the sample is normalised against the absorbance of the well(s) containing blank media
- fluorescence of the sample is normalised using a fitted autofluorescence background curve of the well(s) containing non-fluorescent negative control cells
- the conversion factors generated from calibrant dilutions (Figure 2a) are used to convert GFP fluorescence to MEFL and absorbance to particle count.

**Figure 2.**
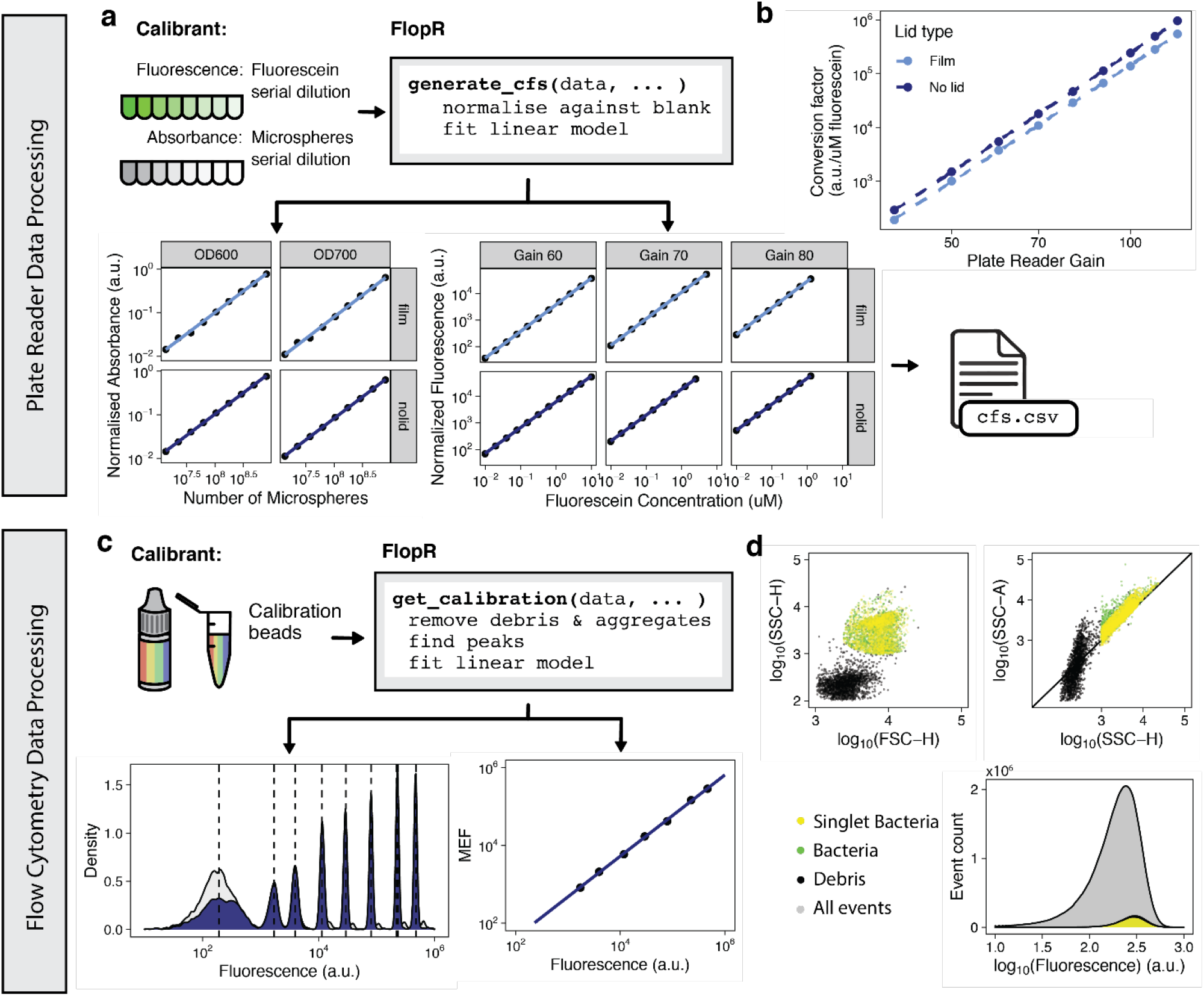
Plate reader and flow cytometry data calibration and processing. (a) FlopR fits plate reader measurements of serial dilutions of fluorescence and absorbance calibrants (fluorescein and microspheres respectively) to generate conversion factors. (b) Conversion factors for fluorescence follow a log linear relationship with gain. (c) FlopR flow cytometry calibration fits measurements of calibration beads, and (d) trims raw flow data to remove debris and identify singlet bacteria.

To generate plate reader conversion factors, a calibration plate should be prepared and measured prior to the main experiment. The calibration plate contains serial dilutions of both plate reader calibrants, and its fluorescence and absorbance are measured under all the potential experimental conditions (e.g. plate type, well volume, lid type) and with a range of gain settings (protocol adapted from (1) available in the supplemental). FlopR takes this data as input, fits a linear model to it on a log-log scale and produces plots showing the model fits for each measured condition. Representative examples of these are shown in Figure 2a. Once a calibration plate has been measured, the conversion factors generated from the data (`cfs.csv`) can be used for all future experiments using the measured conditions, and only needs to be repeated if the plate reader performance is thought to have changed over time. The calculated fluorescence conversion factors and gain of the plate reader follow a log linear relationship, shown in Figure 2b. FlopR fits this relationship to get the final required conversion factor for any specific experimental gain, reducing error of conversion factors of gains that were directly measured.

To process flow cytometry data:

- bacterial clusters are identified using a mixture model from the flowClust package (6)
- singlets are identified using the singletGate function from flowStats (7), as shown in Figure 2d
- peaks from the calibration beads are identified using kernel density estimation and a calibration curve is produced to convert sample data units to Molecules of Equivalent Fluorophore (MEF), which for the case of green fluorescence are units of MEFL.

We use a kernel density clustering model, as opposed to Gaussian mixtures models or k-means clustering models used in other flow cytometry data calibration software (8,9), as it requires less tuning to identify closely clustered peaks (shown in Figure S1). MEFL fluorescence values for the specific bead lot or brand used need to be input by the user.

### Normalisation of autofluorescence in bulk plate reader data

Careful normalization of plate reader data is required to correctly remove autofluorescence from the media and cells themselves. Incorrect normalization can diminish measured fluorescent output dynamic range, particularly in the case of GFP (5). Often normalisation is done simply by subtracting the measured fluorescence of a negative control well containing non-fluorescent cells from the fluorescence of the cells of interest at the same timepoint. This time-based normalisation does not take into account growth differences between the two types of cells, which is common when cells contain genetic circuits that can affect growth (10). Another approach involves normalizing using spectral unmixing of the main identified autofluorescent agents (11, 5); however, this approach requires detailed knowledge of the source of the autofluorescence (5). A better approach is to fit an absorbance versus fluorescence background calibration curve of the negative control cells (Figure 3a) to get fluorescence as a function of absorbance for normalisation (12–15). FlopR has the options to fit a generalized additive model (GAM), locally estimated scatterplot smoothing (LOESS), second-order polynomial or exponential model to the data. This relationship is monotonic from lag phase to early stationary phase, however during stationary phase, we have observed that fluorescence can increase while absorbance remains stable. Furthermore, over very long time periods, certain negative wells showed a decrease in absorbance whilst maintaining the same level of fluorescence (shown in Figure S2), which may be due to changes in cell physiology or stressors. Both of these conditions can lead to reduction in the accuracy of the model fitting. FlopR outputs a plot of the background autofluorescence fit, which should be checked to ensure correct fit. If necessary, alternative negative control wells can be selected, or the data can be trimmed to before the physiologically stressed cell state.

**Figure 3.**
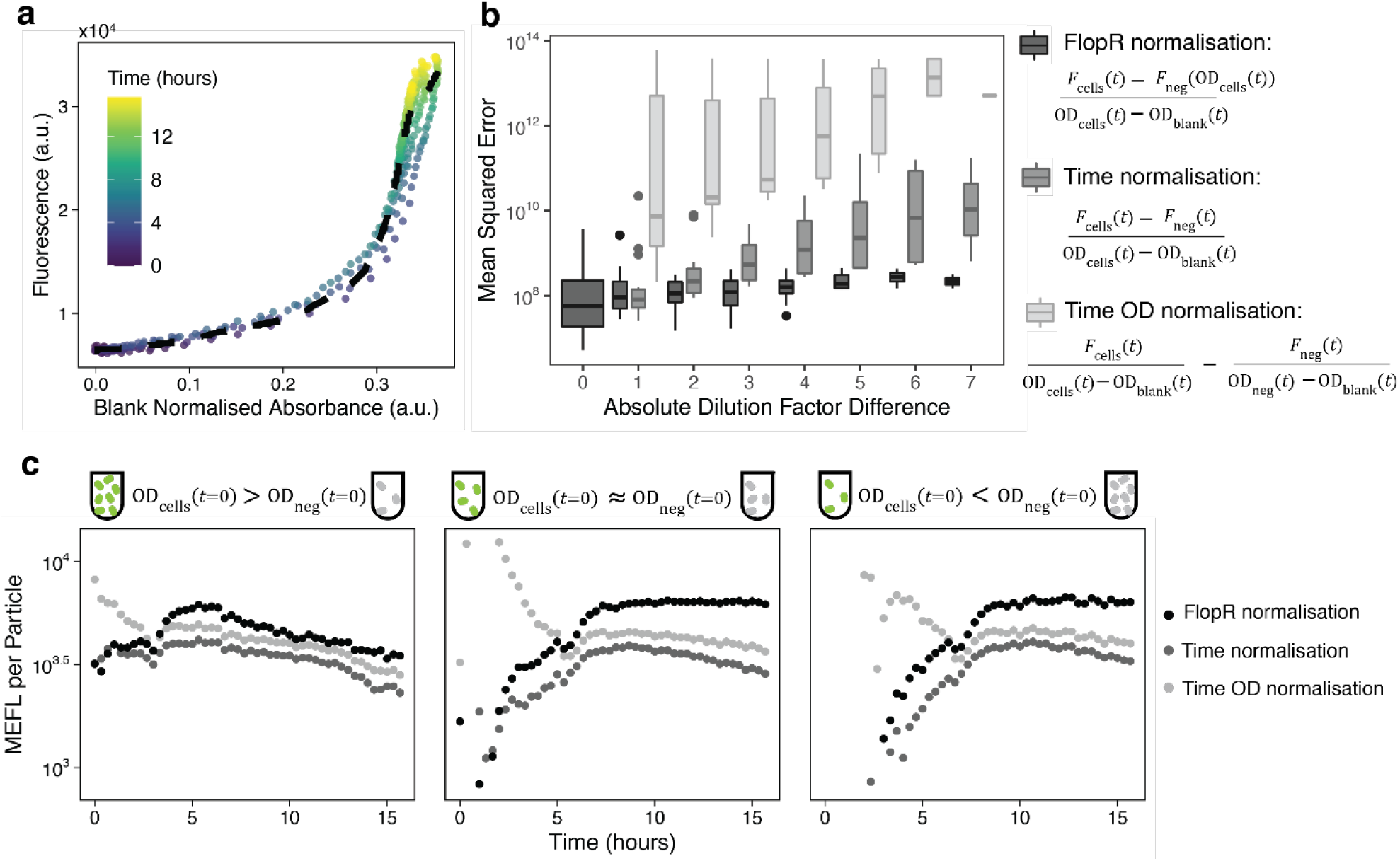
Fluorescence normalization in plate readers using fitted negative control cell autofluorescence. (a) LOESS fit of the absorbance vs fluorescence of negative control cells used for FlopR normalisation. (b) Comparison of normalization performance for negative control cells using FlopR normalisation, time normalisation, or time OD normalisation. Cells were serially diluted by a factor of 3:8, the absolute dilution factor difference is the number of dilution steps between the normalising and normalised cells. (c) MEFL per particle timecourses of single wells calculated using time normalisation, time OD normalisation or FlopR normalisation at different starting concentration of fluorescent and negative non fluorescent control cells.

In order to show quantitatively the performance of the different normalization approaches in cases of growth differences between fluorescent cells and their negative controls, we grew different starting dilutions of both cell types for 15 h (Figure S3). Figure 3b shows the mean squared error of fluorescence per OD of each negative well normalized using the other negative wells at differing starting dilutions. Accurate normalization should remove all autofluorescence, so the ‘ground-truth’ fluorescence is expected to be zero. As the difference in dilution increases, the deviation from zero of both time normalization methods increases substantially (normalisation plots of individual wells are shown in Figure S4). FlopR fitted normalization also shows a moderate increase, however, the increase is considerably smaller than the other types of normalization (Figure 3b is plotted on a log-scale), indicating more consistent normalization performance, regardless of growth. This is demonstrated in Figure 3c, which shows MEFL per particle of fluorescent wells normalized for cases in which the negative wells were at a different starting cell densities. Time-based normalization is particularly prone to poor normalization at early time points, when delays or differences in the start of exponential growth of the two populations will be emphasized.

### Applying FlopR to constitutive GFP expression data

We then sought to demonstrate the use of FlopR using constructs that constitutively expressed GFP with differing strength promoters (Figure 4a). Figure 4b shows the results of measurements of three different strength constructs on a plate reader and flow cytometer, verified against literature values of the same constructs (1). We followed the protocol conducted by the iGEM interlab study, so it is worthwhile to note that these were normalized following the interlab protocol and were not normalized for autofluorescence according to FlopR’s normalization fit. In this case the negative control cells and GFP-expressing cells showed similar growth dynamics, so time-based normalization is expected to give accurate results.

**Figure 4.**
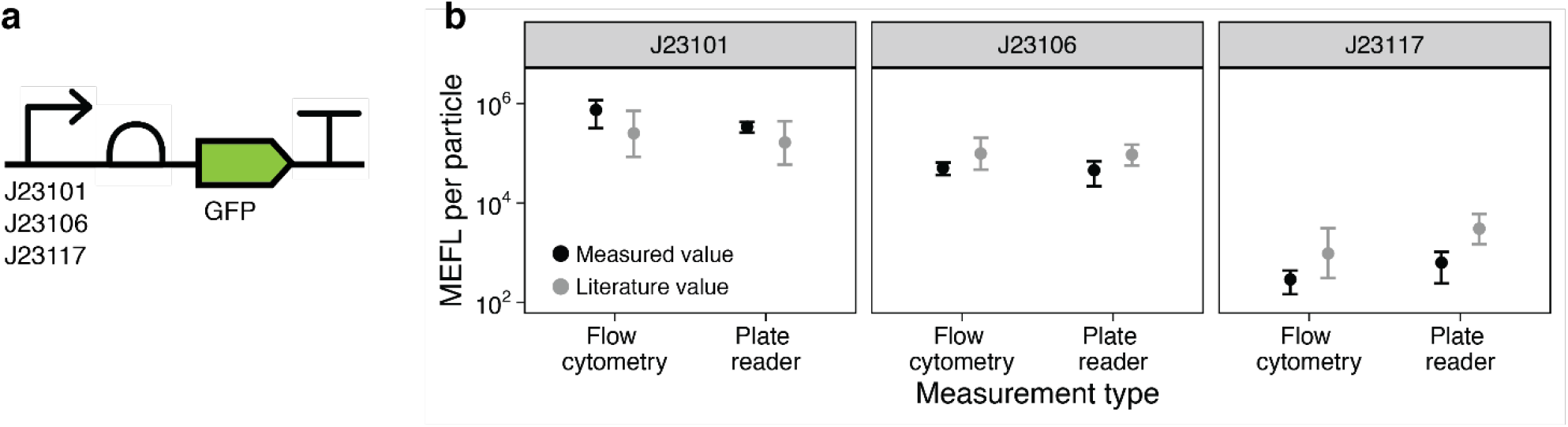
Demonstration of FLOPR-produced MEFL per particle values measured by flow cytometry or on a plate reader. (a) Three different constitutive GFP expression constructs were tested, with promoters J23101, J23106 and J23117. (b) Fluorescence measurements on different days compared to literature values for the same plasmids (1) plate reader data is mean and standard deviation of 12 replicates on 2 days, flow cytometry data is the mean and standard deviation of the median log fluorescence intensities of 11 replicates over 2 days.

### Identifying bacterial subpopulations in bulk plate reader measurements

Finally, we used FlopR to identify specific subpopulations of cells in mixed cultures. There has been growing interest in engineering multicellular consortia (16), however identifying individual strains’ growth dynamics within a mixed population remains challenging. We use a previously described system (17) in which bacteriocin-producing fluorescent ‘killer’ cells were mixed at different starting fractions with non-fluorescent competitor cells, and use FlopR to investigate their interaction dynamics from data measured in a microplate reader and flow cytometer over 12 hours (Figure 5a). Traditionally, separation of the two population fractions is only possible by clustering in flow cytometry data, as shown in Figure 5b. In this case, population fractions are simply calculated by taking the ratio of the cells in the two population clusters. The killer cells were engineered to produce two fluorescent proteins, GFP and red fluorescent protein (RFP), so that clustering could be performed using both fluorescence channels without the issue of spectral overlap of the competitor and killer cell populations when fluorescence expression is weak. To reconstruct population fractions from plate reader data, a positive control well composed of only fluorescent killer cells was measured and used to create an absorbance-fluorescence curve (Figure S5). The measured absorbance of mixed culture wells was then used to get the expected fluorescence value for an 100% killer cell culture, and the ratio of actual measured fluorescence to expected fluorescence taken to obtain the fraction of killer cells present in the mixed culture. This fraction was then multiplied with the calibrated particle count of the mixed wells to get the growth timecourses of individual subpopulations, shown in the top panels of Figure 5d. At high starting concentrations, the killer cells immediately overwhelm the competitor cells and dominate the culture (last panel Figure 5d). However, at low initial killer cell concentrations, the competitor cells experience an initial growth phase, until the concentration of bacteriocin is sufficiently large to begin eliminating the competitors. The calculated fraction of killer cells from measurements in both instruments are shown in the bottom panel of Figure 5d, and a comparison is presented in Figure 5c, both showing that FlopR is able to closely reconstruct accurate subpopulations of cells from bulk data. Reconstruction of cell population fractions from plate reader measurements does not require spectral separation of the two subpopulations and therefore the use of both fluorescent protein channels is not necessary. The reconstruction cell fractions using the green fluorescence channel is shown in Figure S6.

**Figure 5.**
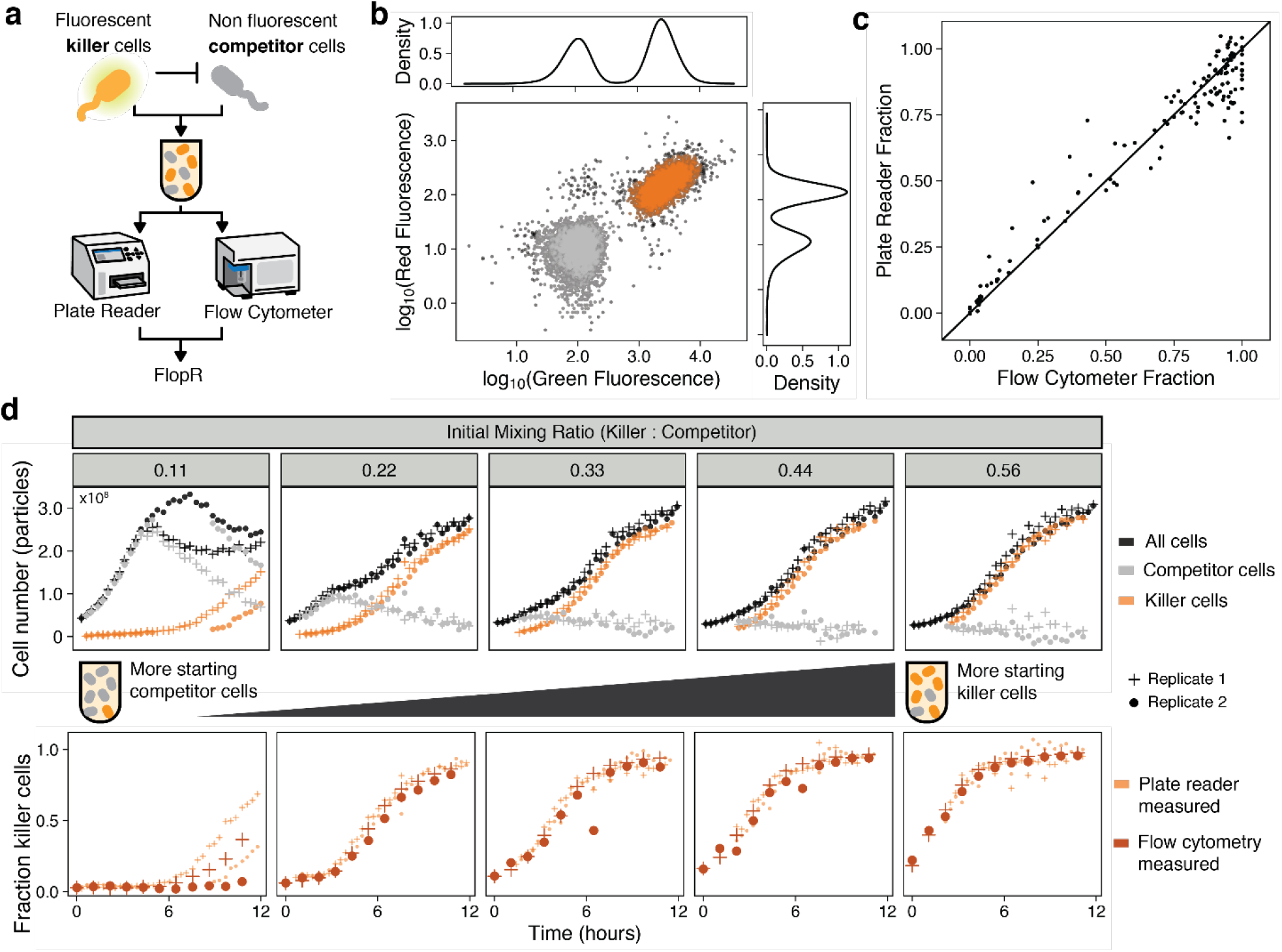
Using FlopR to monitor population dynamics of individual populations within heterogeneous bulk plate reader measurements. (a) Mixed co-cultures of fluorescent killer cells and non-fluorescent competitor cells at different starting ratios were measured and compared using FlopR. (b) Distinct subpopulations are easily identifiable by clustering flow cytometry data. (c) Comparison of plate reader calculated killer cell fractions and flow cytometry measured killer cell population fractions for wells at identical timepoints. (d) dynamics of individual cell populations (orange or grey for killer and competitor respectively) reconstructed from plate reader data using FlopR. The top panel shows timecourses of cell number of strains reconstructed from plate reader data. The bottom panel shows timecourse data of killer cell population fractions calculated from plate reader data or measured from clustered flow cytometry data. Data for (d) is shown from two replicate wells measured on the same day.

## Discussion

FlopR is a free and open source software tool that allows direct comparison of flow cytometry and plate reader green fluorescence data. Future extensions to the software should include fluorescence calibration for colours other than green fluorescence. Possible calibrants for red fluorescence, such as TexasRed, are currently being investigated. Size calibration (18) – currently available in other flow cytometry packages (19) – can also be added, which may be of particular use in order to understand changes or differences in morphology of different bacterial strains. Size calibration may be particularly important to understand which size of microsphere to use as a calibrant to avoid over- or under-estimation of population size, as differences in particle size can cause large differences in light scattering, particularly when the size of the particle approaches that of the incident light wave, as is the case with bacteria (2). Alternatively, the cells themselves could be used to create the calibration curves for specific strains: by measuring the number of cells in a known dilution in the flow cytometer, it would be possible to back calculate the number of cells in a dilution series of the same sample measured in the plate reader. In addition, determination of the MEFL per individual GFP protein, for example by flow cytometry and liquid chromatography-mass spectrometry (20), would be useful to gain quantitative information of how many proteins per cells are producing that fluorescence. This would be highly beneficial to enable the coupling of computational models to experimental data. As this software is open source, it can also be built upon for added functionalities and calibration of other instruments and methods, such as fluorescence microscopy (21), or other imaging techniques.

Measurement standards help reproducibility, transparency and collaboration within the scientific process. Despite this, fluorescence remains identified as a problematic measurement area (22,23). We hope this tool will be of use to the synthetic biology community to facilitate and encourage the use of standardized units in fluorescence measurements. The full details on how to use the package can be found in the READme.txt file on FlopR’s GitHub repository.

## Methods

### Instruments and settings

All plate reader experiments were done on a Tecan SPARK Multimode Microplate Reader, using 96 well black, clear bottom plates (Corning). Specific plate reader settings for the individual experiments of each figure are outlined in Tables S1. All fluorescence measurements were made from the top of the plate, with 125 μL cell cultures. Plate reader data for Figures 3 and 5 was collected throughout the timecourse. Plate reader data for Figure 4b was collected at a single time point, following the protocol detailed in Beal *et al*. (1).

All flow cytometry experiments were carried out on an Attune NxT Flow Cytometer. Green fluorescence was measured on the BL1 channel (excitation laser: 488 nm, emission filter: 530/30 nm), and red fluorescence on the YL2 channel (excitation laser: 561 nm, emission: 620/15 nm). Cell cultures were diluted 1 μL in 200 μL of filtered PBS, with 3 mixing cycles and minimum 3 wash rinses between samples.

Plate reader laser excitation and emission wavelengths for experiments in Figures 3 and 5 differ by a couple of nm from the flow cytometer excitation and emission wavelengths of the BL1 channel. However, as both the plate reader’s excitation and emission bandwidths are 20 nm and the flow cytometer bandpass filter is 30 nm, the small difference in the settings is negligible, and the measurements from both instruments are still comparable.

### Calibration materials

The full protocol for generating serial dilutions of plate reader calibrants is detailed in method S1. Briefly, fluorescein (Sigma-Aldrich) dissolved in PBS and 0.89um monodisperse silica microspheres (Cospheric) suspended in microbiology grade water were used to create serial dilutions in a 96 well black, clear bottom plate (Corning), with four replicate dilution series for each calibrant. Due to the quick settling time of the microspheres, their dilutions were remixed by pipetting up and down immediately before measurement on the plate reader. For flow cytometry calibration, one drop of Spherotec Rainbow calibration particles (catalog number: 422903, lot number: 073112) was mixed with 300 μL of PBS and measured before, and at the same settings, as the relevant experiment.

### Strains & cultures conditions

All plasmids, strains, and antibiotic working concentrations used are listed in Table S1. For Figure 3 and 5 cells were cultured in M9 minimal media supplemented with 0.4% glycerol and 0.2% casamino acids throughout the experiments. Cells for the experiment in Figure 4b were cultured in LB media throughout the experiment, following the protocol in (1). For Figures 3, overnight cultures were diluted 1:1000 in fresh M9 media, grown for 6 more hours at 200 rpm, diluted to a target OD 700 nm of 0.1, then diluted by a factor of 3/8 (for initial starting dilution ratios of: 1, 0.375, 0.141, 0.053, 0.020, 0.007, 0.003 and 0.001). For Figure 5, overnight cultures were diluted 1:100, then mixed at the specified ratios.

## Author Information

### Author Contributions

AJHF and CPB conceived the idea. CR, AJHF and KYW performed all experiments. CR and AJHF developed the software. CR and AJHF performed analysis of the results. CR and AJHF wrote the first draft of the manuscript. All authors contributed to manuscript revision, read and approved the submitted version.

### Notes

The authors declare no competing financial interest.

## Acknowledgments

The authors would like to thank Dr Stefanie Frank (UCL) for providing plasmids from the iGEM biobrick registry plate distribution.

