## Supplementary Information for "FlopR: An open source software package for calibration and normalization of plate reader and flow cytometry data"

### Supplemental method 1:
### Microsphere & Fluorescein Plate Reader Calibration Protocol

This protocol is used to generate conversion factors for the conversion of plate reader data from fluorescence arbitrary units per absorbance to units of Molecules of Equivalent Fluorescein (MEFL) per particle. Once converted, the plate reader data can be compared to calibrated flow cytometry data.

Fluorescein is used as a calibrant to generate fluorescence conversion factors (MEFL/a.u.), and 0.890um diameter microspheres (Cospheric) are used to generate absorbance conversion factors (particles/Abs).

The dilutions outlined below are to create a calibration plate for 125 µL cultures, and are compatible with the example calibration_plate_layout.csv. The concentrations of calibrants in the plate layout file can be changed by the user if necessary.

**Materials:**

- 10ul 2.5mM fluorescein stock solution (stored in the fridge, light sensitive).
- 7ml of PBS
- 0.1g of 0.89um diameter microspheres (stored in small glass vial with green taped lid).
- 10ml of molecular biology grade water
- 96 well clear bottom black plate
- Breathable film

**Part 1: Fluorescein and Microsphere dilutions.**

**Fluorescein dilution:** Rows A-D in the plate. This is a 1:2 dilution.

1. Pipette 250 µl of 10 µM fluorescein into wells A1, B1, C1 and D1
2. Pipette 125 µl of PBS into wells A2-A12, B2-B12, C2-C12 and D2-D12 (it is easiest to do this using the multichannel pipette and a sterile trough).
3. Using the multichannel pipette, pipette 125 µl of fluorescein from wells A1-D1 into A2-D2, and mix by pipetting up and down.
4. Continue the serial dilutions, 125 µl of A2 into A3, 125 µl of A3 into A4 ect… DO NOT pipette any fluorescein into the wells in column 12 (these wells serve as PBS blanks).

**Microspheres dilution:** Rows E-H on the plate. This is a 3:5 dilution done in 1.5 ml Eppendorf tubes and then pipetted into the plate (modified from the 1:2 dilution from the iGEM interlab and Beal et al 2019). This was done to avoid saturation issues that were observed with larger dilution steps, and to allow vortexing/good mixing of the tubes to minimize error due to microsphere settling (see note below).

Note: Microspheres settle very fast when diluted in a liquid, make sure to mix/vortex well before any steps.

1. Create a microsphere stock solution with 0.1 g of microspheres and 1.3 ml of deionized water
2. Make 500 µl of a second stock solution by adding 300 µl of the above microsphere solution to 200 µl of water.
3. Make 1 ml of a third stock solution by diluting the second stock 1:10 (100 µl of microsphere to 900 µl of water)

4. Prepare eleven 1.5 ml Eppendorf tubes (numbered 1 to 11) with each containing 600 µl of water.
5. Add 900 µl of the 2nd stock of microspheres to Eppendorf tube 1 (and mix well by pipetting up and down)
6. Add 900 µl from tube 1 to tube 2 and mix well
7. Add 900 µl from tube 2 to tube 3 and mix well
8. … continue the serial dilution until tube 11. Tubes 1-10 should have 600 µl of diluted microspheres, tube 11 should have 1500 µl.
9. Pipette 125 µl of tube 1 into wells E1, F1, G1 and H1
10. Pipette 125 µl of tube 2 into wells E2, F2, G2 and H2
11. … continue until tube 11 into wells E11, F11, G11 and H11
12. Pipette 125 µl of molecular biology grade water into wells E12, F12, G12 and H12.

**Plate reader measurement:**

1. Measure the calibration plate on the plate reader using all intended experimental settings:
    - Excitation and emission settings identical to the green fluorescence channel of the flow cytometer that the plate reader data will be compared to.
    - Gain in steps of 10, from 40-120
    - With and without a film
2. Save both files as csvs (ex. Calibration_YYYY_MM_DD_film.csv and Calibration_YYYY_MM_DD_nofilm.csv).

**Part 2: Generating all cfs using calibration data:**

1. Install the FlopR library, to install FlopR run devtools::install_github("ucl-cssb/flopr")
2. To generate conversion factors run:
   flopr::generate_cfs(calibration_dir, date, microsphere_vals = c(4,12))
    - calibration_dir is the directory in which the calibration data and calibration plate layout is stored, it should be named as the date on which the calibration was done (ex. 20200601 for 1st of June 2020). The film measurements should be named film.csv, no film measurements should be named nolid.csv, the calibration plate layout is called calibration_plate_layout.csv.
    - Date should be input as YYYYMMDD
    - microsphere_vals is the values of the microsphere concentrations that should be used to fit, by default FlopR uses the highest 8 values (values 4 to 12), this can be modified by the user if saturation issues occur.

The function will output plots of absorbance conversion factors and fluorescence conversion factors for each measured condition, and a file of the conversion factors called cfs_generated.csv, which has a column names cf (the conversion factor), and slope. If the calibration has been performed correctly the slope should be close to 1.

**Table S1:** Plate reader settings for all figures and experiments.

| Figure | Gain | Absorbance | Excitation<br>Bandwidth: 20 nm | Emission<br>Bandwidth: 20 nm | Shaking | Lid-Type |
|---|---|---|---|---|---|---|
| 3 | 135 | 700 nm | 485 nm | 535 nm | 150 rpm continuous double orbital shaking, 2 mm amplitude | Breathable film |
| 4b | 80 | 600 nm | 488 nm | 530 nm | Continuous shaking following protocol in (1) | No lid |
| 5 | 125 | 700 nm | 485 nm (GFP)<br>561 nm (mCherry) | 530 nm (GFP)<br>620 nm (mCherry) | 150 rpm continuous double orbital shaking, 2 mm amplitude | No lid |

**Table S2:** Plasmids, strains and antibiotic and their working concentrations used in this study.

| Plasmid | Strain | Antibiotic | Relevant Figure | Description | Source |
|---|---|---|---|---|---|
| none | MG_Gm_CFP | Gentamicin (10 µg/mL) | Fig 3a, b<br>Fig 5d | Non green fluorescent negative control cells, competitor cells | (17) |
| pMPES_AF01,<br>P63_AF043 | JW3910 | Gentamicin (10 µg/mL) | Fig 3c,<br>Fig 5d | Positive control fluorescent cells, Bacteriocin producing | (17)<br>Keio collection *metB* KO |
| BBa_J364000 | NEB DH5alpha | Chloramphenicol (25 µg/mL) | Fig 4b, first panel | Strong constitutive green fluorescent cells | iGEM biobrick registry plate distribution |
| BBa_J364001 | NEB DH5alpha | Chloramphenicol (25 µg/mL) | Fig 4b, second panel | Medium constitutive green fluorescent cells | iGEM biobrick registry plate distribution |
| BBa_J364002 | NEB DH5alpha | Chloramphenicol (25 µg/mL) | Fig 4b, third panel | Weak constitutive green fluorescent cells | iGEM biobrick registry plate distribution |

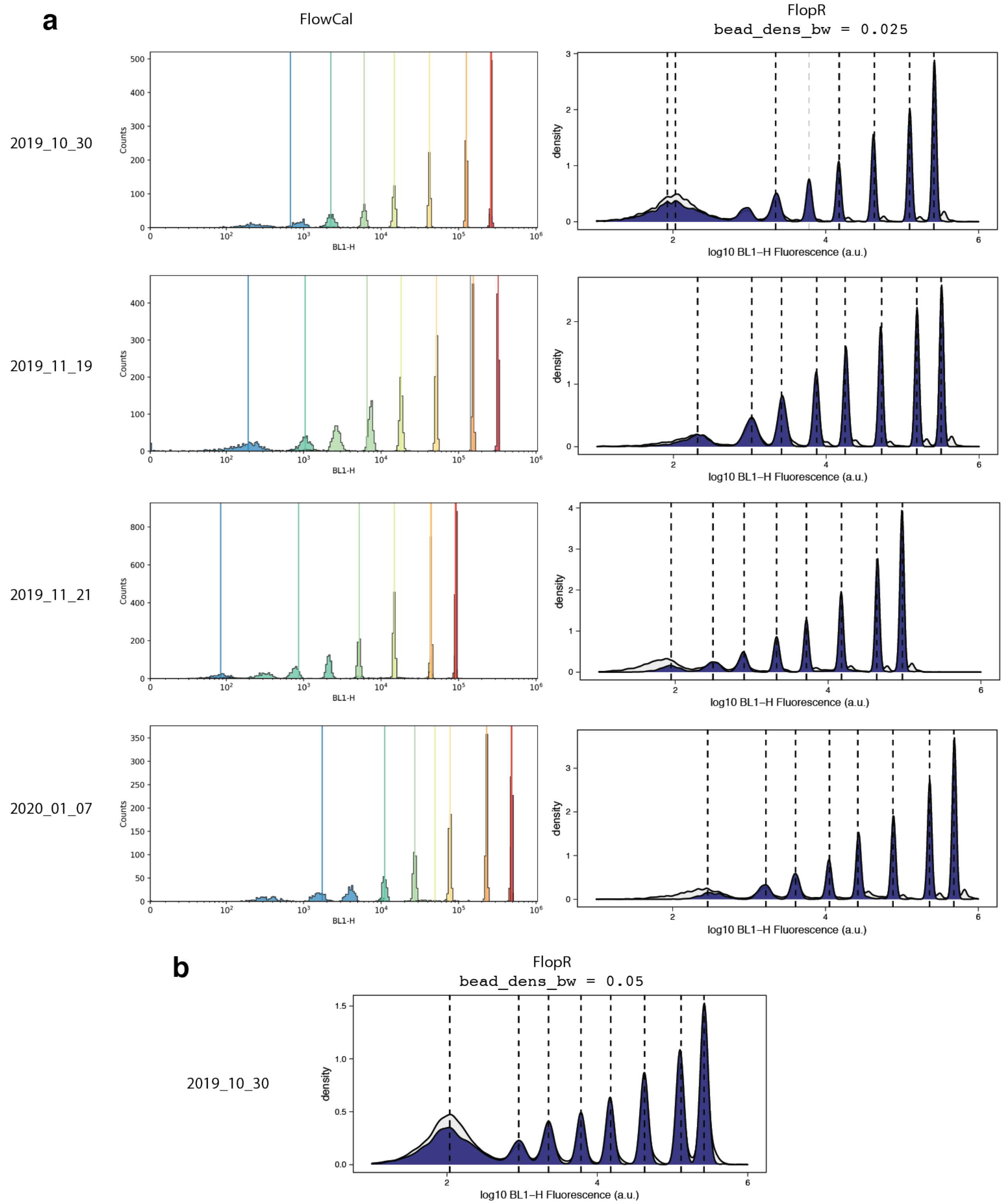


**Figure S1:** (a) Identification of peaks from a gaussian mixture (GM) model (left column) in the FlowCal package vs FlopR's peak identification using kernel density estimation (right column). The GM model incorrectly identifies peaks in bead data measured on four different dates. FlopR's kernel density estimation peak identification identifies peaks correctly in 3 out of 4 cases at FlopR's default kernel density bandwith parameter (set by the `Bead_dens_bw` parameter). `Bead_dens_bw` can be specified by the user if erroneous peaks are being found, the default value is 0.025, (b) changing the parameter to 0.05 gives correct bead identification for case 1 (2019_10_30).

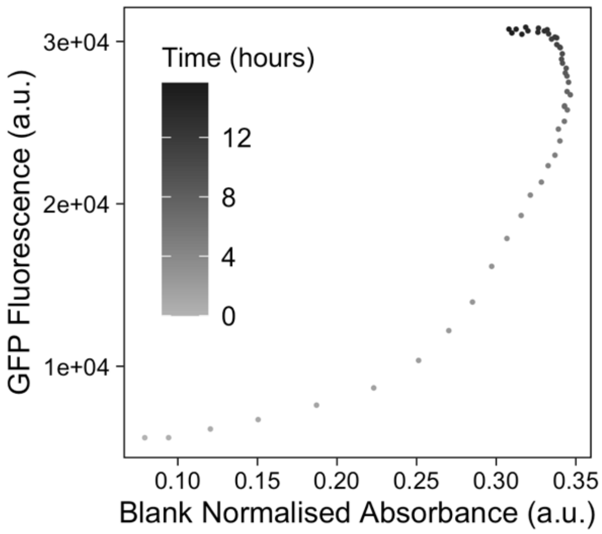


**Figure S2:** A negative control well from the experiment shown in Figure 3 that shows a decrease in absorbance while maintaining the same level of fluorescence at late timepoints. This may be due to cell stressors or change in the cells' physiological state when in late stationary phase (>12 hours).

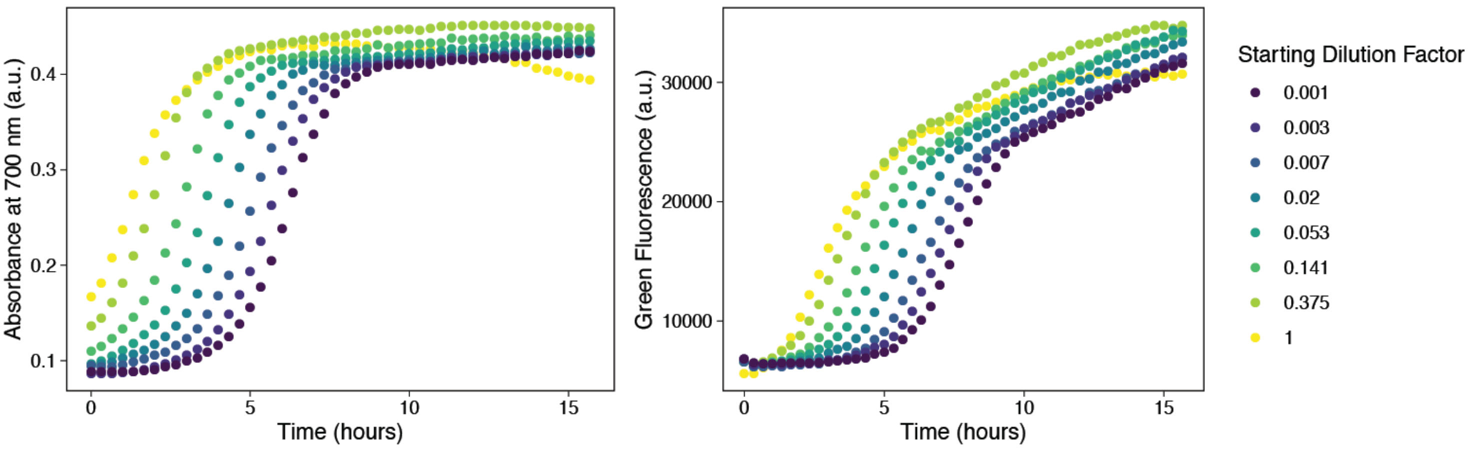


**Figure S3**: Raw absorbance (700nm) and green fluorescence measurements of the experiment shown in Figure 3. Overnight cultures were diluted to different starting concentrations to result in staggered growth.

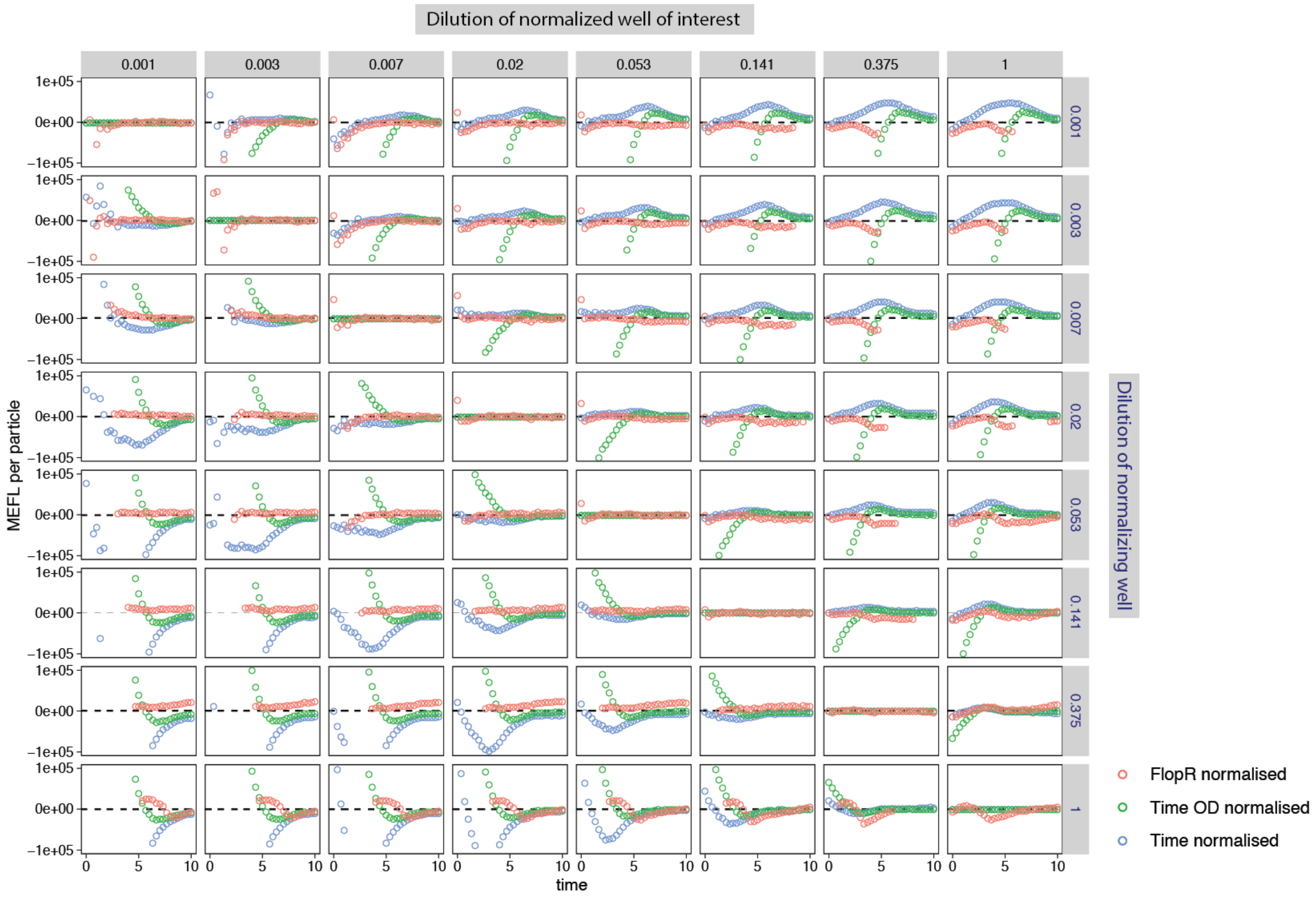


**Figure S4**: Negative control wells normalized (using a LOESS fit) for autofluorescence (ideal normalization should produce a timecourse that remains at 0 MEFL per particle). The starting dilutions of the normalized wells are listed along the top of the plot, and starting dilutions of normalizing wells are listed on the right (plots on the upper right show the result of normalisation with a normalising well with a lower starting dilution than the well of interest, plots of the lower left shown the result of normalisation with a normalising well with a higher starting dilution than the well of interest). Missing points in FlopR normalisation are due to the fact that LOESS models do not extrapolate the fit

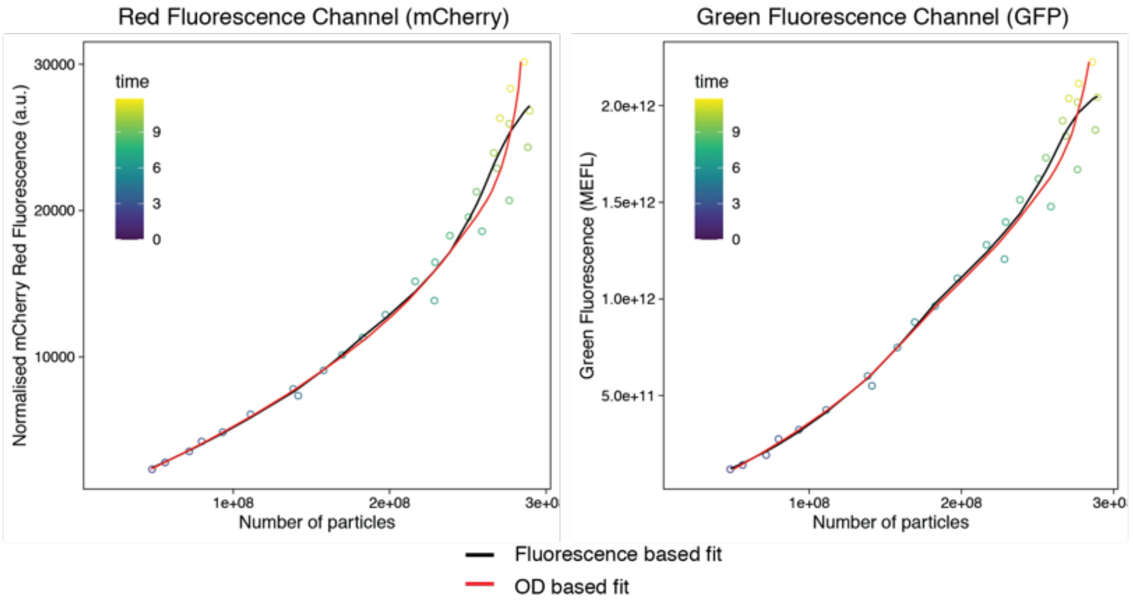


**Figure S5**: Fluorescence vs absorbance curve of the positive control, only killer cell wells, used to calculate population fractions from plate reader data. The left panel is the fraction calculation curve used for Figure 5 (mCherry red fluorescence), however population fraction calculation is also possible using green fluorescence as the killer cells are also constitutively expressing GFP (right panel). The black curve shows models fit based on fluorescence, the red curve is models fit to number of particles.

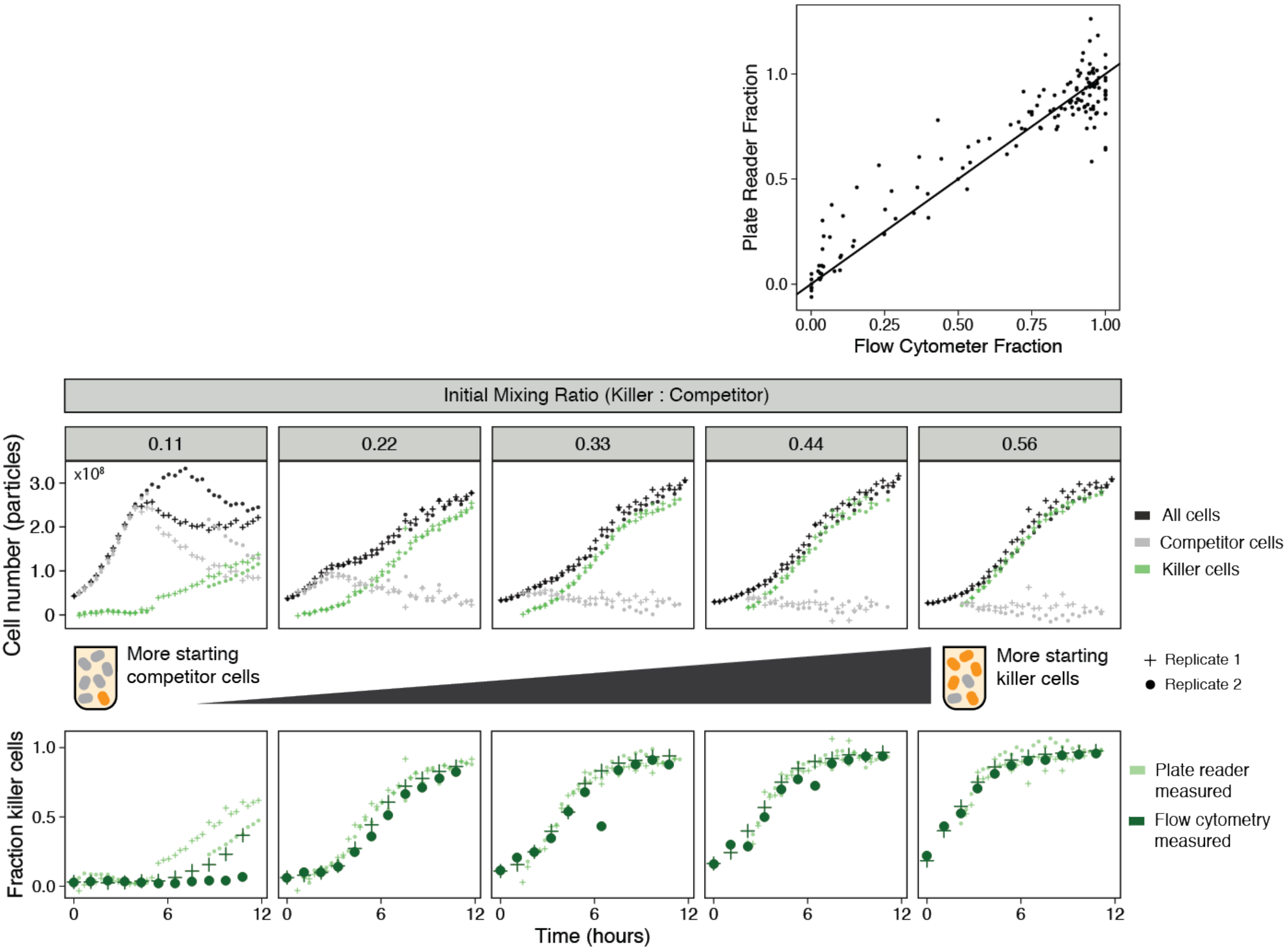


**Figure S6**: Using GFP fluorescence fraction calculation curve (Figure S5, right panel) to recreate subpopulation fractions. Calculating population fractions using green fluorescence gives a similar output to using red fluorescence (as is shown in Figure 5)
